# Myonuclear alterations associated with exercise are independent of age in humans

**DOI:** 10.1101/2022.09.20.506578

**Authors:** E. Battey, J.A Ross, A. Hoang, D.G.S. Wilson, Y. Han, Y. Levy, R.D. Pollock, M. Kalakoutis, J.N. Pugh, G.L. Close, G. M. Ellison-Hughes, N.R. Lazarus, T. Iskratsch, S.D.R. Harridge, J. Ochala, M.J. Stroud

**Affiliations:** Centre for Human & Applied Physiological Sciences, Faculty of Life Sciences & Medicine, King’s College London, London, UK; British Heart Foundation Centre of Research Excellence, School of Cardiovascular Medicine and Sciences, King’s College London, London, UK; Novo Nordisk Foundation Center for Basic Metabolic Research, University of Copenhagen, Copenhagen, Denmark; School of Engineering and Materials Science, Queen Mary University of London, London, UK; Randall Centre for Cell and Molecular Biophysics, Faculty of Life Sciences & Medicine, King’s College London, London, UK; School of Sport and Exercise Sciences, Tom Reilly Building, Byrom Street, Liverpool John Moores University, Liverpool, UK; Department of Biomedical Sciences, Faculty of Health and Medical Sciences, University of Copenhagen, Copenhagen, Denmark

**Keywords:** exercise, ageing, nuclei, nuclear shape, nuclear lamina

## Abstract

Age-related decline in skeletal muscle structure and function can be mitigated by regular exercise. However, the precise mechanisms that govern this are not fully understood. The nucleus plays an active role in translating forces into biochemical signals (mechanotransduction), with nuclear lamina protein Lamin A regulating nuclear shape, nuclear mechanics, and ultimately gene expression. Defective Lamin A expression causes muscle pathologies and premature ageing syndromes, but the roles of nuclear structure and function in physiological ageing and in exercise adaptations remain obscure. Here, we isolated single muscle fibres and carried out detailed morphological and functional analyses on myonuclei from young and older exercise-trained individuals. Strikingly, myonuclei from trained individuals were more spherical, less deformable, and contained a thicker nuclear lamina than untrained individuals. Complementary to this, exercise resulted in increased levels of Lamin A and increased myonuclear stiffness in mice. We conclude that exercise is associated with myonuclear remodelling, independently of age, which may contribute to the preservative effects of exercise on muscle function throughout the lifespan.

**Key points:**

- The nucleus plays an active role in translating forces into biochemical signals
- Myonuclear aberrations in a group of muscular dystrophies called laminopathies suggest that the shape and mechanical properties of myonuclei are important for maintaining muscle function.
- Here, we present striking differences in myonuclear shape and mechanics associated with exercise, in both young and old humans.
- Myonuclei from trained individuals were more spherical, less deformable, and contained a thicker nuclear lamina than untrained individuals.
- We conclude that exercise is associated with age-independent myonuclear remodelling, which may help to maintain muscle function throughout the lifespan.

## Introduction

Human lifespan has increased substantially over the past half-century and this trend is projected to continue (UN, 2022). However, this has not been accompanied by an equivalent extension of the healthspan in old age; instead, morbidity has been extended, and independence and quality of life attenuated (Brown, 2015). Thus, a ‘managed compression of morbidity’ is essential to address social and economic issues associated with an extended lifespan (Brown, 2015). A contributing factor to morbidity is the decline in skeletal muscle structure and function associated with ageing. Muscle contractions produce force, allowing us to carry out whole-body movements such as walking, stair-climbing or rising from a chair – movements essential for independence and quality of life. The ageing process, however, is compounded by the physically inactive status of individuals in a technologically advanced society (Guthold *et al*., 2018; Nikitara *et al*., 2021). Furthermore, physical inactivity can accelerate the decline in physiological function that inevitably occurs during later years of life (Lazarus & Harridge, 2017; Shur *et al*., 2021).

Skeletal muscle structure and function can be better maintained in old age by exercise, but the mechanisms behind this remain poorly understood (Wroblewski *et al*., 2011; Pollock *et al*.,2015; Lazarus *et al*., 2019). An area of research which is understudied is how exercise and ageing influence the ability of skeletal muscle to translate force into biochemical signals (mechanotransduction) at the subcellular level. This is pertinent given the contractile nature of muscle and the opposing effects of exercise and inactivity on the frequency and intensity of muscle contractions. Emerging data suggest that the nucleus is a critical mechanosensor that orchestrates cell structure, function, and adaptive responses (Kirby & Lammerding, 2018). Indeed, the shape and mechanical properties of nuclei appear to regulate gene expression by altering genome organisation and ultimately influencing broader transcriptional profiles (Tajik *et al*., 2016; Kirby & Lammerding, 2018; Piccus & Brayson, 2020; Kalukula *et al*., 2022). Additionally, altered nuclear shape and nuclear envelope stretching can expand nuclear pore complexes and ion channels, facilitating translocation of mechanosensitive transcription factors Yes-associated protein/Transcriptional coactivator with PDZ-binding motif (Yap/Taz) and myocardin-related transcription factor (MRTF-A) or ions such as Ca^2+^, respectively, altering gene expression and signalling (Kirby & Lammerding, 2018; Maurer & Lammerding, 2019; Ross & Stroud, 2021; Shen *et al*., 2022).

Abnormal nuclear structure and responses to forces are hallmarks of numerous diseases that result in skeletal muscle weakness and premature ageing, commonly caused by mutations in nuclear envelope and associated proteins (Goldman *et al*., 2004; Ross *et al*., 2019; Battey *et al*., 2020; Earle *et al*., 2020; Kalukula *et al*., 2022). Such proteins physically link the nucleus to the cytoskeleton, providing a nexus for mechanotransduction (Crisp *et al*., 2006; Banerjee *et al*., 2014; Kirby & Lammerding, 2018; Ross & Stroud, 2021). The nuclear lamina, which lines the inner nuclear membrane and comprises Lamins A/C, B1 and B2, tethers chromatin to the nuclear periphery, associates with nuclear pore complexes, and connects the nucleoskeleton to the cytoskeleton via Linker of Nucleoskeleton and Cytoskeleton (LINC) complex (Osmanagic-Myers *et al*., 2015; Stroud *et al*., 2017; Stroud, 2018; Owens *et al*.,2021). Within this prominent location, the nuclear lamina is critically positioned to sense cytoskeletal forces to regulate gene expression, biochemical signalling and overall cell function and adaptation (Cho *et al*., 2017; Maurer & Lammerding, 2019).

Importantly, various diseases caused by mutations in genes encoding nuclear lamina proteins (termed laminopathies) primarily affect mechanically active muscle tissue and result in aberrant nuclear shape, structural integrity and mechanotransduction (Janin *et al*., 2017; Earle *et al*., 2020; Shin & Worman, 2021). One such laminopathy is Hutchinson-Gilford Progerin (HGPS) syndrome, a premature ageing syndrome caused by a mutation in the gene encoding Lamin A/C (Goldman *et al*., 2004; Merideth *et al*., 2008). Defective Lamin A/C expression also results in muscular dystrophy characterised by muscle weakness (such as autosomal dominant Emery-Dreifuss muscular dystrophy, limb-girdle muscular dystrophy type1B, and *Łmna*-congenital muscular dystrophy) (Bonne *et al*., 1999; Maggi *et al*., 2016). Thus, myonuclear shape, nuclear envelope proteins, and nuclear mechanics are dysregulated in muscle pathologies and premature ageing syndromes and may have important roles in age-related muscle dysfunction.

Lamin A/C has been shown to be required for normal nuclear mechanics in myotubes and for cardiac and skeletal muscle overload hypertrophy responses in mice (Cupesi *et al*., 2010; Earle *et al*., 2020; Owens *et al*., 2021). Indeed, a congenital mutation in Lamin A/C causing muscular dystrophy resulted in altered nuclear mechanics, attenuated hypertrophy and force capacity in response to functional overload in mouse skeletal muscle (Owens *et al*., 2021). In cardiac tissue, in response to pressure overload, haploinsufficient Lamin A/C mice demonstrated reduced ventricular mass and myocyte size and impaired mechanotransduction (Gerhart-Hines *et al*., 2007; Little *et al*., 2010; Gurd, 2011). Collectively, these studies hint at a potential Lamin A/C-dependent mechanosensitive signalling cascade in regulating both muscle hypertrophy and oxidative exercise adaptations.

Despite the large amount of evidence suggesting the importance of nuclear shape, mechanics and lamina in premature ageing and muscle pathologies, their roles in normal ageing and exercise are poorly understood (Gerhart-Hines *et al*., 2007; Little *et al*., 2010; Cupesi *et al*., 2010; Gurd, 2011; Earle *et al*., 2020; Owens *et al*., 2021). To this end, we investigated whether ageing and exercise affected structure and function of skeletal muscle nuclei. Single muscle fibres from young and older trained and untrained individuals were isolated and myonuclear structure and function analysed. Detailed 2D and 3D morphological analyses of myonuclei revealed striking nuclear shape differences in trained individuals compared to untrained individuals, regardless of age. Additionally, myonuclei from trained individuals had increased nuclear lamina deposition and were less deformable compared to untrained counterparts. Consistently, skeletal muscle from trained mice had increased levels of Lamin A and increased nuclear stiffness. Our data suggest for the first time in humans that exercise is associated with differences in myonuclear shape and mechanics, that likely mitigate the deleterious effects of inactive ageing.

## Results

### Myonuclei in both younger and older trained individuals are more spherical compared to untrained counterparts

Nuclei from laminopathy patients with premature ageing and muscle dysfunction are known to be ruffled and elongated (Goldman *et al*., 2004; Park *et al*., 2009; Tan *et al*., 2015; Earle *et al*., 2020). Here, we hypothesised that myonuclei would show similar abnormalities in physiological ageing. To investigate the effects of age and exercise training on myonuclear shape, muscle fibres were isolated from vastus lateralis biopsies taken from younger untrained (YU, 33 ± 9.5 years), younger trained (YT, 32 ± 5.4 years), older untrained (OU, 79 ± 11.3 years), and older trained (OT, 76 ± 3 years old) individuals.

In contrast to our hypothesis, myonuclei were strikingly rounder in shape in both younger and older trained individuals (Figure 1A), consistent with a previous report in mice (Murach *et al*., 2020). Indeed, the aspect ratio of myonuclei from trained individuals was ~27-29% lower than untrained counterparts, demonstrating significant differences in roundness (YU, 2.4 ± 0.3; OU, 2.3 ± 0.3; YT, 1.7 ± 0.1; OT, 1.6 ± 0.2; Figure 1B). To control for possible differences in fibre tension that may confound interpretation, we normalised myonuclear aspect ratio to sarcomere length and importantly found no differences associated with sarcomere length (Figure 1C). As further controls, exercise-dependent alterations to myonuclear shape remained apparent in slow muscle fibres expressing Myosin Heavy Chain 7, indicating that fibre-type differences between groups did not influence the effects observed (Figure 1D). In contrast to changes in nuclear aspect ratio, nuclear area was comparable in muscle fibres from trained and untrained individuals, highlighting that differences to myonuclear aspect ratio were driven by in myonuclear shape rather than size (Figure 1E).

**Figure 1:**
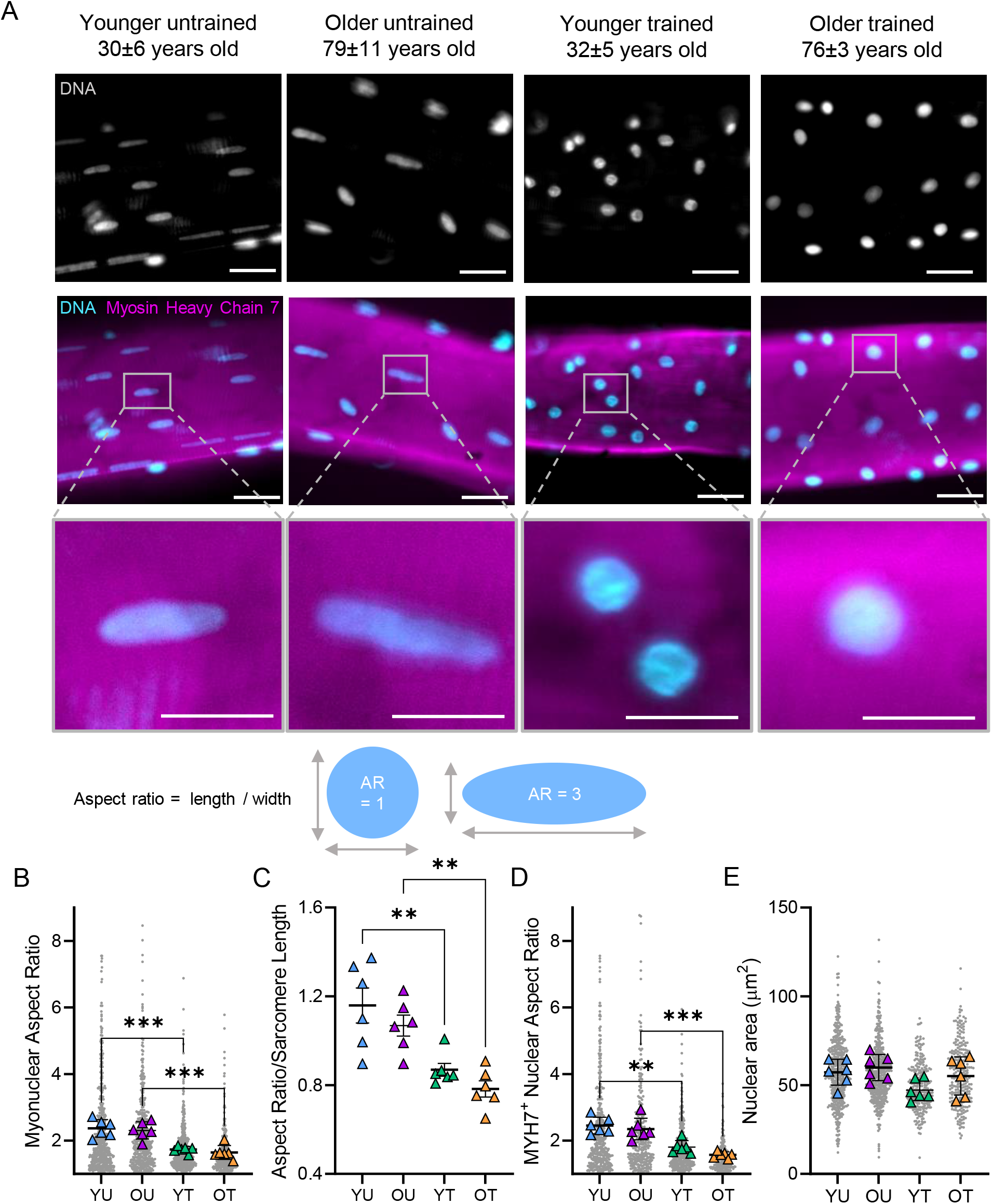
Altered 2D myonuclear shape in trained younger and older individuals. (A) Representative images of vastus lateralis muscle fibres isolated from younger untrained (YU), older untrained (OU), younger trained (YT) and older trained (OT) individuals, stained with DAPI (cyan) and Myosin Heavy Chain 7 (magenta) to visualise myonuclei and slow myosin, respectively. Scale bars 30 μm and 10 μm main images and zoomed insets of myonuclei, respectively. (B-E) Calculation of aspect ratio (length/width of nucleus) and nuclear area (μm^2^) shown above graphs. (B) Comparisons of myonuclear aspect ratio in YU, OU, YT, and OT individuals; 2053 total nuclei analysed (C) Comparisons of myonuclear aspect ratio between groups after normalisation to sarcomere length (D) Myonuclear aspect ratio in MYH7+ fibres (1385 total nuclei analysed) (E) Comparisons of nuclear area (μm^2^) between groups (1453 total nuclei analysed). (B-E) Coloured symbols represent individual means, unfilled grey symbols represent myonuclei; mean values for individuals were used for two-way ANOVA tests (n = 6, ** P < 0.01 *** P < 0.001); error bars represent mean ± SD.

Next, we performed 3D shape analysis of myonuclei by acquiring serial optical z-slices through whole muscle fibres (Figure 2A-B). In line with 2D shape changes observed, myonuclei from OT displayed a significant reduction in 3D aspect ratio compared to OU, with a trending reduction observed in YT compared to YU (P < 0.07) (Figure 2A). Furthermore, sphericity values were higher in trained individuals compared to age-matched untrained counterparts, indicating nuclei were more spherical in these groups (Figure 2B). Importantly, nuclear volumes were comparable across groups, consistent with our 2D analyses (Figure 2C).

**Figure 2:**
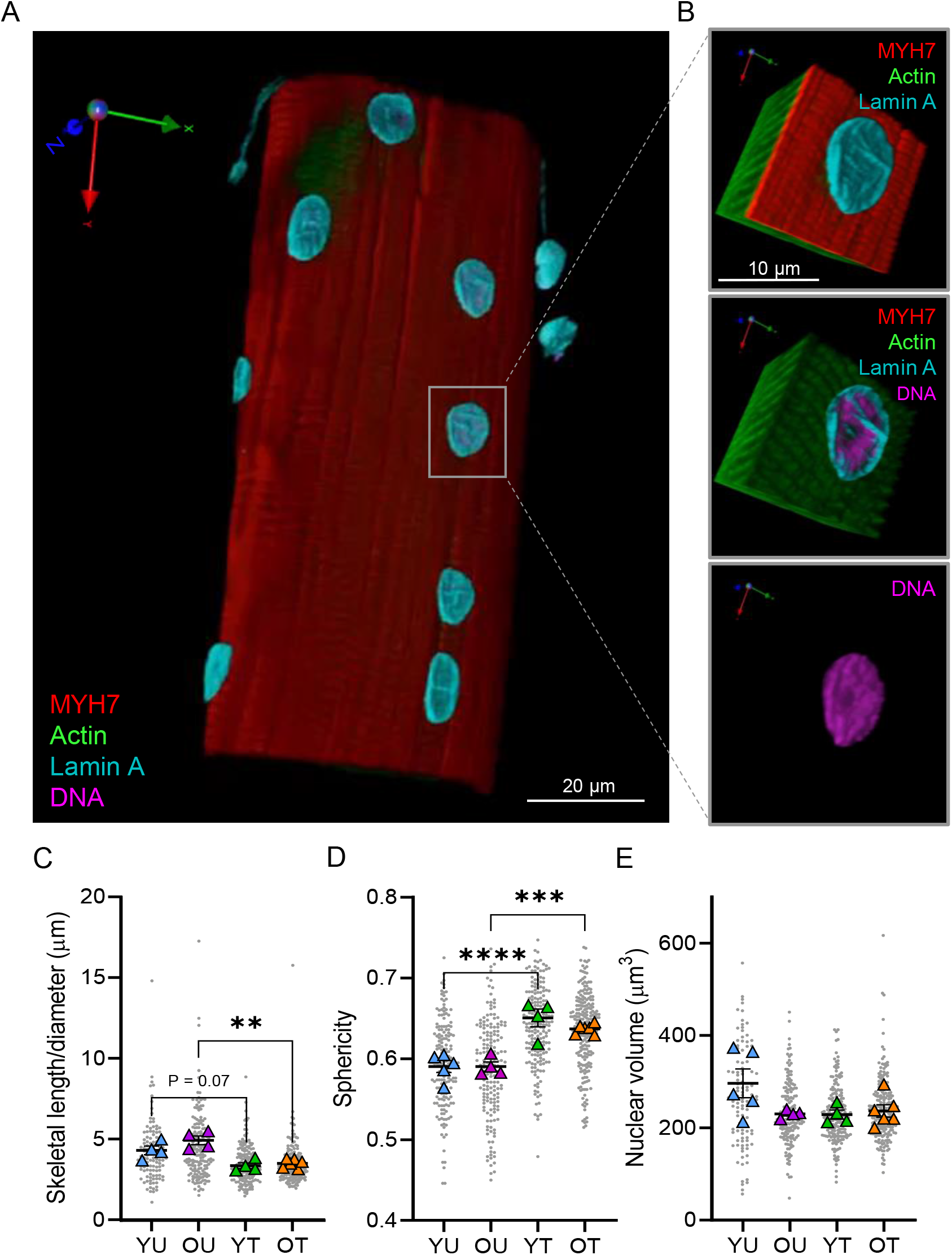
Lower 3D aspect ratio and greater sphericity in myonuclei from trained younger and older individuals. (A) Representative three-dimensional rendering of z-stack images of a single human vastus lateralis muscle fibre acquired with a spinning disk confocal microscope equipped with a 63X oil objective lens. Muscle fibre stained to visualise Lamin A (cyan), DNA (magenta), Actin (green), Myosin Heavy Chain 7 (MYH7, red). (B) Representative zoomed images of 3D-rendered nucleus. (C-E) Comparisons of nuclear skeletal length/ diameter (μm), sphericity, and volume in younger untrained (YU), older untrained (OU), younger trained (YT) and older trained (OT) individuals. Coloured symbols represent individual means, unfilled grey symbols represent myonuclei; mean values for individuals were used for two-way ANOVA tests (n = 5-6, ** P < 0.01 *** P < 0.001 **** P < 0.0001); error bars represent mean ± SD.

Taken together, 2D and 3D analyses of myonuclear shape revealed striking morphological differences in younger and older trained individuals compared to untrained counterparts.

### Nuclear lamina deposition is greater in skeletal muscle fibres from trained individuals

Lamin A localisation and levels regulate nuclear stiffness and nuclear roundness (Lammerding *et al*., 2006; Swift *et al*., 2013; Earle *et al*., 2020; Srivastava *et al*., 2021). Additionally, it has recently been shown that a *Lmna* congenital muscular dystrophy alters mechanotransduction in cultured myotubes and attenuates the hypertrophic response to functional overload in mouse skeletal muscle *in vivo*, implicating a role of Lamin A/C in exercise adaptations (Owens *et al*., 2021). Thus, to investigate whether exercise affects the organisation of Lamin A in skeletal muscle fibres from YU, YT, OU and OT individuals were stained with a Lamin A-specific antibody (Figure 3). As expected, Lamin A localised to the periphery of myonuclei (Figure 3A). Importantly, we observed a significant increase in nuclear lamina deposition in myonuclei from OT compared to OU (Figure 4B-C). Next, we quantified nuclear invaginations, which are tube-like infoldings of the nuclear envelope and are reported to play roles in premature ageing syndromes (McClintock *et al*., 2006; Frost, 2016; Schoen *et al*., 2017). However, our data showed there were no significant differences in total invagination length (μm) between myonuclei from all groups (Figure 4D-E).

**Figure 3:**
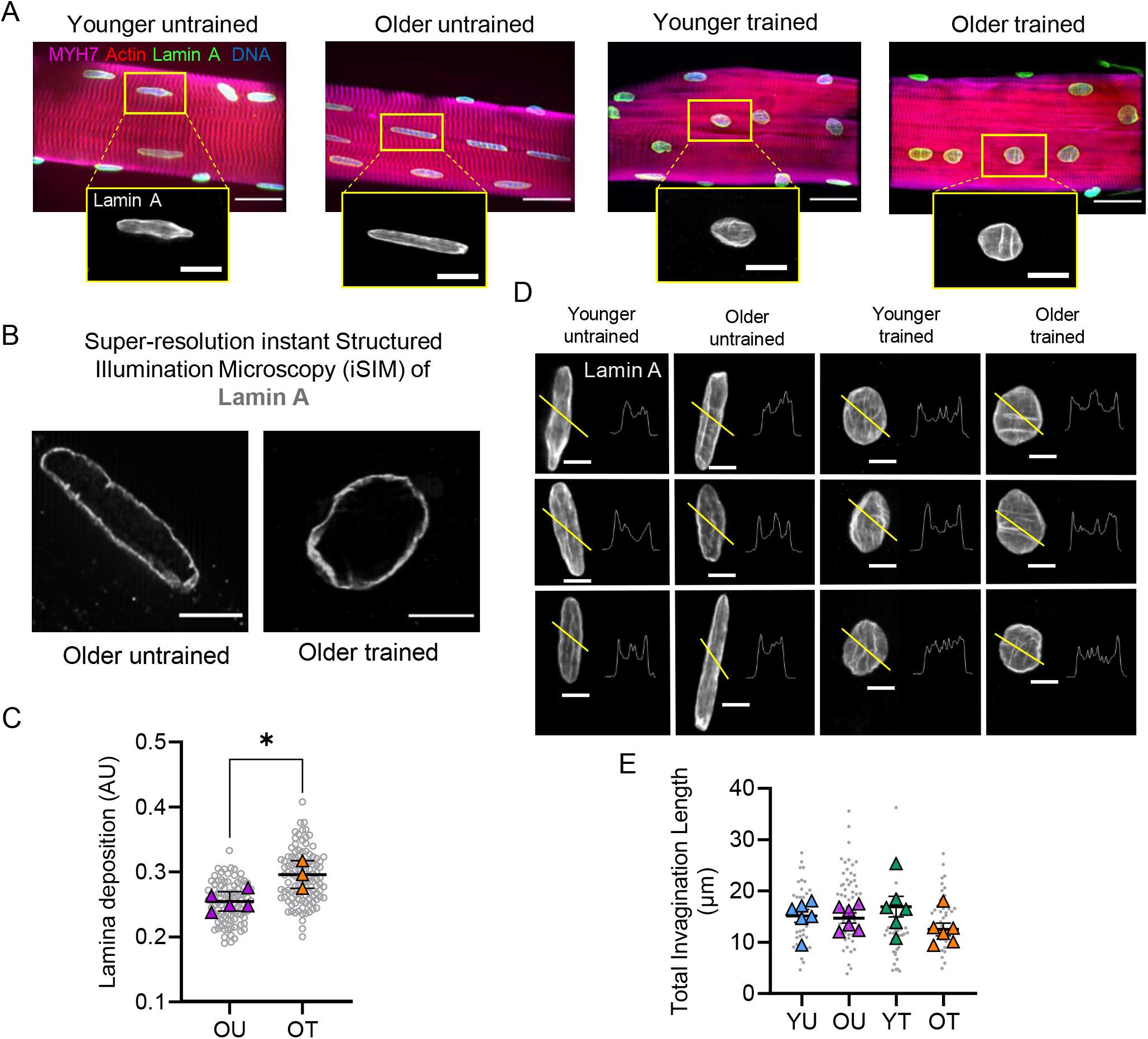
Organisation of Lamin A in trained individuals and untrained counterparts. (A) Representative images, acquired through confocal microscopy using a 63x oil objective, of muscle fibres isolated from younger untrained (YU), older untrained (OU) patients, younger trained (YT) and older untrained (OT). Fibres were stained with DAPI to visualise DNA (blue), Actin (red), Lamin A (green, gray) and Myosin Heavy Chain 7 (MYH7, magenta). Scale bar 25 μm in main images, 10 μm in zoomed images. (B) Representative images of myonuclei from OU and OT muscle fibres acquired through super resolution iSIM microscopy. Scale bars 5 μm. (C) Quantification of Lamin A deposition (μm) in muscle fibres from OT and OU. n = 3-4 per group, unpaired t-test revealed significant difference between groups (P < 0.05). (D) Standard deviation projections of Lamin A-stained myonuclei and pixel intensity line scans (yellow line) from YU, OU, YT, and OT. (E) Lamin A total invagination length (μm) in muscle fibres from YU, OU, YT, OT, n = 6. Two-way ANOVA revealed no significant differences between groups. Coloured symbols represent individual means, grey symbols represent myonuclei; mean values for individuals were used for two-way ANOVA tests and t-test. Error bars represent mean ± SD.

**Figure 4:**
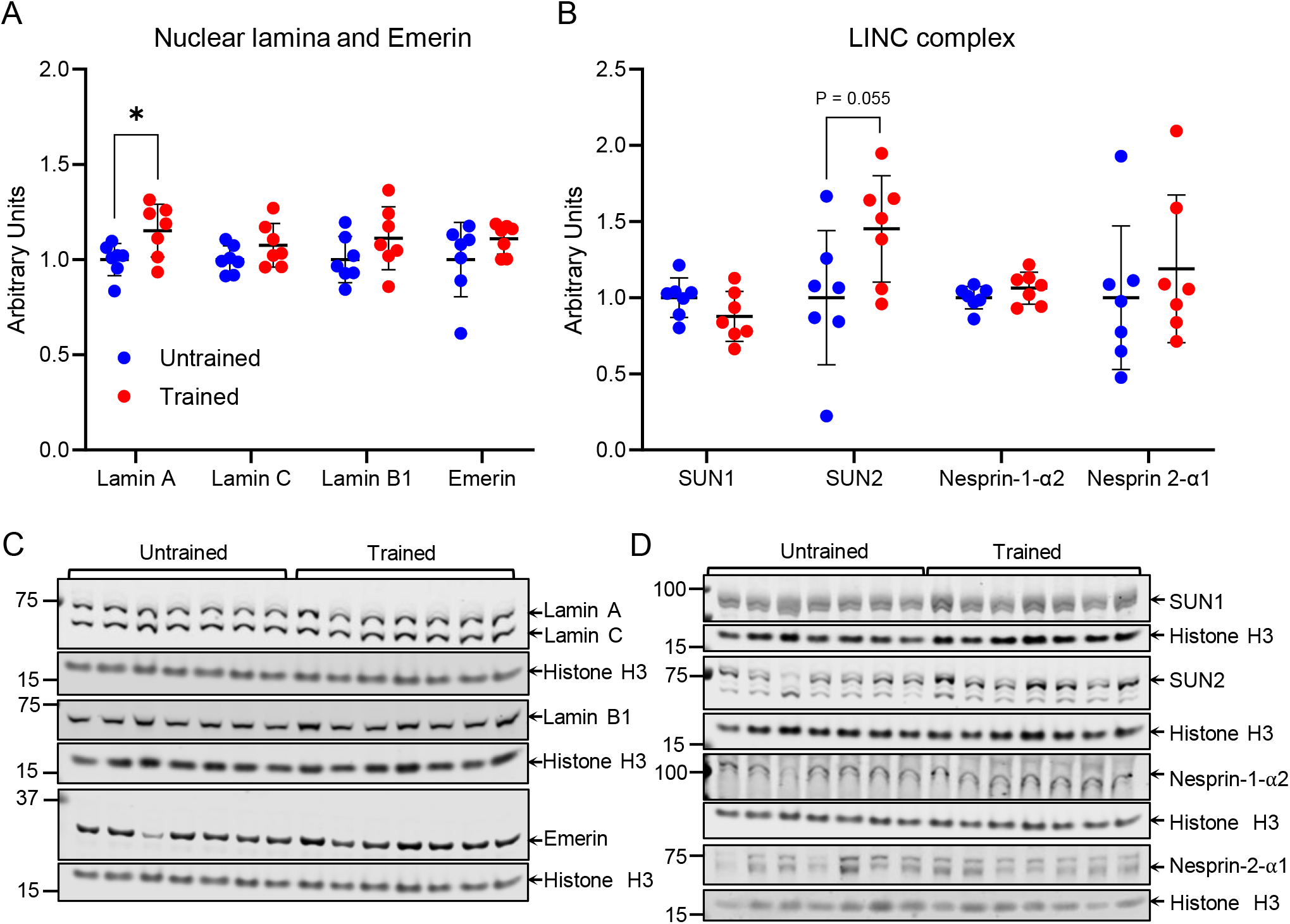
Lamin A levels are increased in trained mouse tibialis anterior muscle. (A) Protein levels of Lamin A, Lamin C, Lamin B1 and Emerin normalised to Histone H3 in tibialis anterior muscle from untrained and high intensity endurance trained mice (B) Protein levels of Linker of Nucleoskeleton and Cytoskeleton (LINC) complex proteins SUN1, SUN2, Nesprin-1-α2, and Nesprin-2-α1 normalised to Histone H3 in tibialis anterior muscle from untrained and trained mice (C-D) Images of western blots from which data in A-B were obtained. Note that Lamin A levels were significantly increased and SUN2 levels trending to increase. Arrows indicate predicted molecular weights (kD). Data points represent individual mice, n = 7 per group, * indicates P < 0.05 (t-test). Error bars represent mean ± SD.

These data suggest that training is associated with increased nuclear lamina deposition in primary human skeletal muscle.

### Lamin A levels are increased in exercise-trained mice

Given the precious nature and paucity of human muscle biopsy samples for protein and biophysical analysis, next we used a mouse model to investigate the effects of exercise on myonuclear parameters further.

To determine whether exercise affected the protein levels of nuclear lamins and LINC complex proteins, we performed western blotting on tibialis anterior muscle tissue from mice following 8 weeks of treadmill running. In line with increased Lamin A deposition in trained human muscle fibres, Lamin A levels were significantly increased in trained mice compared to untrained counterparts (442 ± 53 vs 384 ± 33, respectively, P < 0.05; Figure 4A). In contrast, levels of Lamins C, B1, and Emerin were not significantly different (Figure 4A). LINC complex proteins, which connect the cytoskeleton to the nucleus via the nuclear lamina, were then analysed. Consistent with the increase in Lamin A, levels of SUN2, which is known to preferentially bind Lamin A over Lamin C (Liang *et al*., 2011), showed a 45% increase in trained mice that was trending towards significance (1129 ± 498 vs 1639 ± 394, respectively, P = 0.055). However, other LINC complex proteins SUN1, Nesprin-1α2 and Nesprin-2α1 were not significantly different in trained compared to untrained mice (Figure 4B).

### Exercise alters myonuclear deformability and stiffness

Changes in nuclear shape and Lamin A expression are associated with altered nuclear mechanosensitivity, with Lamin A being a key regulator of the mechanical stiffness of nuclei (Lammerding *et al*., 2006). Thus, we next addressed whether structural alterations in myonuclei from trained individuals translated to biophysical changes in nuclear mechanics. Whilst it is accepted that adaptations to exercise are at least in part driven by mechanotransduction (Hornberger & Esser, 2004; Kirby, 2019; Attwaters & Hughes, 2022), the role of nuclear mechanics in this context has not previously been investigated.

To assess the effects of exercise on myonuclear function, myonuclear deformability was compared in OU and OT human samples. Single muscle fibres were mounted and stretched to different tensions, fixed, and stained to visualise myonuclei and Z-discs (Figure 5A) (adapted from Chang et al., 2010; Chapman *et al*., 2014). We reasoned that if nuclear aspect ratio increased proportionately with sarcomere length, nuclei were considered compliant with fibre tension; conversely, if nuclear aspect ratio did not scale with sarcomere length, they were considered stiffer. As expected, in both OT and OU fibres, there was a positive relationship between sarcomere length and nuclear aspect ratio (R values were 0.36 and 0.45 for OT and OU, respectively; 211 OT and 260 OU nuclei analysed, respectively; P < 0.05; Figure 5B). However, this relationship was significantly steeper in fibres from OU compared to OT (gradients were 0.79 and 0.29, respectively, P < 0.05; Figure 5B). Additionally, the variance of myonuclear aspect ratio normalised to sarcomere length was significantly higher in OU compared to OT fibres, showing less consistency in myonuclear shape changes with stretching in OU fibres (Figure 5C, P < 0.05).

**Figure 5:**
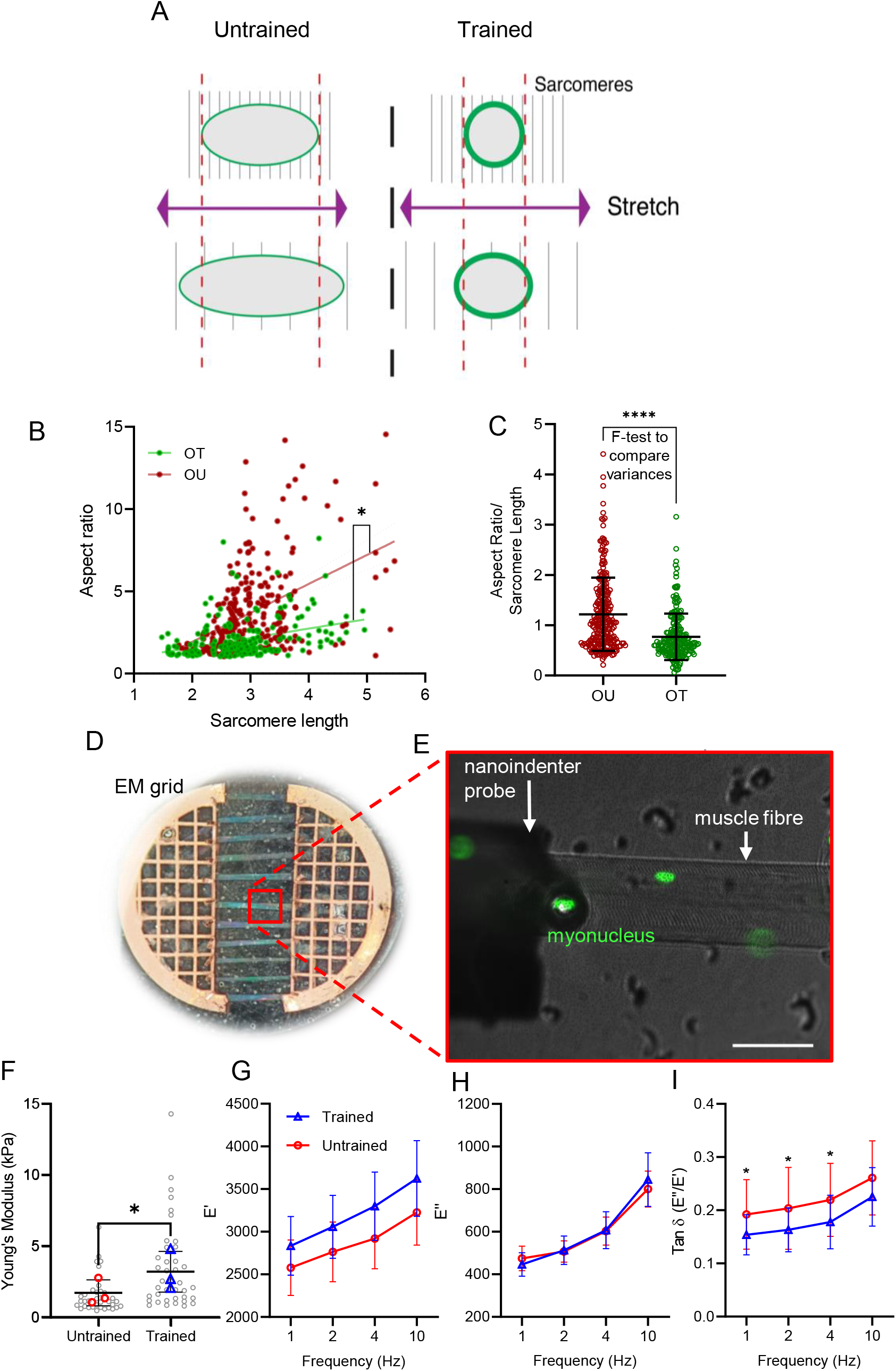
Exercise training results in stiffer myonuclei. (A) Schematic of fibre mounting and stretching. (B) Relationship between sarcomere length and nuclear aspect ratio in muscle fibres from older trained (OT) and untrained (OU) patients. (C) Variance of aspect ratio/ sarcomere length in OT and OU fibres. Note that sarcomere length positively correlates with extent of muscle fibre stretch and that myonuclei in OT fibres were significantly stiffer than myonuclei from OU fibres. * denotes statistically significant difference between gradients of slopes (linear regression analysis, F-test to compare variances). (D) Typical setup of individual muscle fibres after isolation and mounting in parallel on an electron microscopy (EM) grid split in half for imaging and nanoindentation. (E) Nanoindentation of single muscle fibres (brightfield) with myonuclei labelled (green) being probed by nanoindenter (brightfield, left of image). Scale bar, 50 μm. (F) Comparison of Young’s modulus (kPa) in nuclei from untrained (red) and trained mice (blue) (G-I) comparisons of E’, E” and Tan Delta (E”/E’) at different Dynamic Mechanical Analysis (DMA) frequencies (Hz) in nuclei from untrained and exercise trained mice. Note that myonuclei were significantly more stiff and more elastic in fibres from trained vs. untrained mice. Each coloured data point represents the average for each mouse, n = 3 per group. Error bars represent mean ± SEM. * denotes statistically significant difference between groups (t-test and mixed effects analysis).

These data suggest that nuclei in trained individuals were less compliant with increasing fibre tension compared to OU fibres (Figure 5B). In other words, nuclei in fibres from trained individuals appeared stiffer than in untrained individuals. To confirm this, we performed nanoindentation to physically probe nuclei and directly test the effects of exercise on myonuclei (Figure 5D-E). Indeed, we observed an 87% increase in Young’s modulus (kPa) (a measure of stiffness) in myonuclei from trained mice compared to untrained mice (trained 1.7 ± 0.9 vs untrained 3.2 ±1.3, P < 0.05; Figure 5F). Additionally, there was a significant difference in the viscoelasticity, whereby tan δ, the fraction of the loss modulus (the viscous component) over the storage modulus (the elastic component), was on average ~20% lower in exercise trained mice at 1, 2 and 4 Hz DMA (P < 0.05), and 14% lower at 10 Hz DMA (not significant) (Figure 5G-I), indicating more elastic nuclei in the trained mice.

Taken together, analysis of nuclear mechanics in human and mouse muscle fibres indicated that training reduced myonuclear deformability, increased myonuclear stiffness and elasticity.

## Discussion

Muscle pathologies and premature ageing syndromes caused by mutations in nuclear lamina and envelope proteins have revealed a common phenotype: abnormal nuclear shape and defective mechanotransduction. However, whether similar structural and functional defects occur with physiological ageing with and without exercise in human skeletal muscle had not previously been investigated.

Our main findings are that exercise, regardless of age, is associated with more spherical, less deformable myonuclei, with increased Lamin A levels and deposition at the nuclear lamina. This implies that myonuclear mechanotransduction may have a role in governing exercise adaptations (Figure 6A). Maintaining myonuclear structure and function through regular exercise may be an important factor in preserving muscle function throughout the lifespan (Figure 6B). Conversely, myonuclear dysfunction in untrained muscle may contribute to age-related decline in muscle mass and function (Figure 6B). Below, possible mechanisms and consequences of exercise-related and inactivity-related alterations in myonuclear structure and mechanics will be discussed.

**Figure 6:**
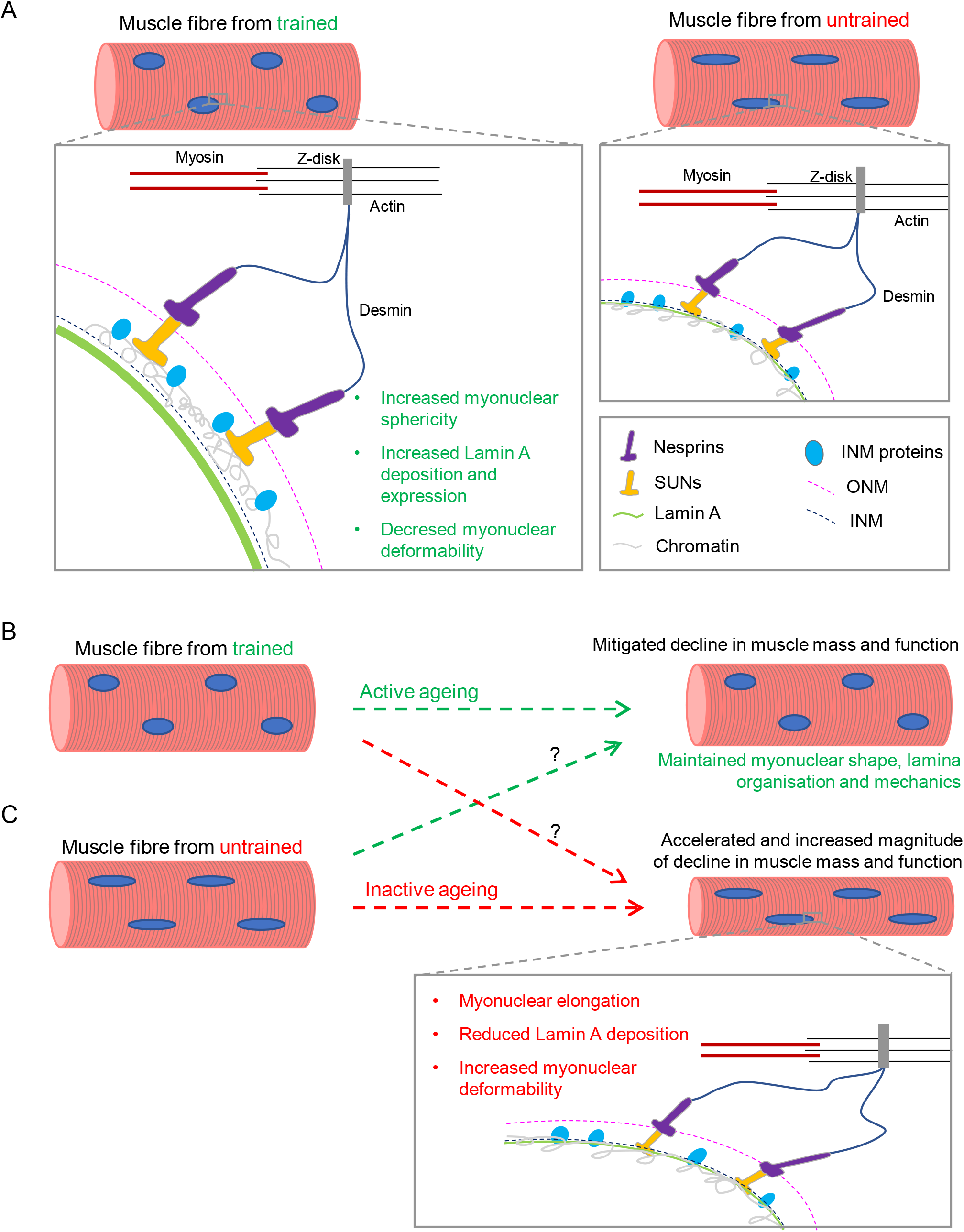
Summary and proposed effects of myonuclear remodelling with training, and inactivity related defects in nuclear mechanotransduction with age. (A) In skeletal muscle fibres from trained individuals, nuclear envelope proteins including the nuclear lamina effectively transduce cytoskeletal forces to the nucleus to regulate signalling pathways. (B) In skeletal muscle from trained older individuals, myonuclear shape and mechanotransduction are preserved (C) In skeletal muscle from untrained older individuals, myonuclei are more elongated, nuclear lamina levels are reduced, and myonuclei more deformable. This may lead to increased susceptibility to myonuclear damage and defective mechanotransduction that results in decline in muscle mass and function.

Greater myonuclear sphericity and stiffness in trained muscle fibres may be a result of the increased Lamin A expression. Lamin A regulates myonuclear shape and mechanics in muscle cells, with loss or misexpression of Lamin A resulting in myonuclear elongation and increased deformability (Roman *et al*., 2017; Earle *et al*., 2020). Forces can also induce conformational changes to nuclear envelope and lamina proteins, modulating the mechanical properties of nuclei (Swift *et al*., 2013; Guilluy *et al*., 2014; Buxboim *et al*., 2014). Thus, increased Lamin A expression may be an initial response to exercise training, which causes increased myonuclear sphericity and stiffness, and reduces myonuclear deformability in trained muscle fibres.

The structural and mechanical alterations to myonuclei in trained individuals may have several consequences beneficial to muscle fibre function (Figure 6). These alterations may be underpinned by altered chromatin organisation and expression of genes important for oxidative capacity or repression of atrophy (Ho *et al*., 2013; Fischer *et al*., 2016). In addition to facilitating exercise adaptations, myonuclear remodelling in trained individuals may be mechanoprotective. Conversely, nuclear defects driven by inactivity may be detrimental for cellular function and health (Kalukula *et al*., 2022). Thus, increased Lamin A expression and stiffness, and reduced deformability of myonuclei in trained muscle fibres may improve resilience against contractile forces during future exercise bouts.

The elongated shape and mechanical properties of myonuclei in untrained individuals were reminiscent of those in muscle fibres from humans and mice with muscular dystrophies characterised by muscle wasting and dysfunction (Tan *et al*., 2015; Earle *et al*., 2020). Thus, defective myonuclear structure and function due to inactivity may contribute to age-related muscle dysfunction. Specifically, chromatin stretching may result in expression of genes that contribute to muscle atrophy, which are elevated after two weeks of inactivity (Jones *et al*.,2004). Additionally, altered chromatin organisation may repress genes encoding contractile or mitochondrial proteins, decreasing force production and endurance capacity (Figure 6B). Deformable myonuclei in untrained individuals may be more susceptible to nuclear envelope rupture, impacting cell health (Earle *et al*., 2020; Kalukula *et al*., 2022). These possible consequences of myonuclear dysfunction in old age may collectively contribute to impairments in muscle mass, strength, and endurance with age, and be alleviated by exercise-mediated myonuclear remodelling (Figure 6B).

A limitation of our work was the limited availability of human muscle biopsies from the four groups, which could be complemented with future studies. In the present investigation, the older untrained group was composed of hip fracture patients with other underlying health conditions (see Table 1) which may have influenced the observed myonuclear aberrations. Participants were mixed sex, and drug administration and variations in habitual dietary intake were not stringently accounted for, possibly introducing variability in muscle fibre size and other variables. To this end, a study group composed of an older untrained group with clearer inclusion and exclusion criteria related to physical activity levels may provide a more accurate representation of the consequences of inactive ageing. Nevertheless, analysis of muscle fibres from this group has provided insight into myonuclear structure and function in elderly inactive individuals. Furthermore, these individuals displayed a strikingly similar nuclear shape phenotype to apparently healthy, younger untrained individuals.

**Table 1.** Participant characteristics.

| Group | Age | Sex | Height (cm) | Weight (kg) | BMI | Comment | Drug intake |
| --- | --- | --- | --- | --- | --- | --- | --- |
| Younger untrained | 25 | F | 164 | 55 | 20.4 |  |  |
|  | 27 | M | 184 | 80 | 23.6 |  |  |
|  | 22 | M | 178 | 75 | 23.7 |  |  |
|  | 42 | F | 172 | 73 | 24.7 |  |  |
|  | 38 | F | 158 | 48 | 19.2 |  |  |
|  | 44 | F | 166 | 52 | 18.9 |  |  |
| Young marathon runners (younger trained) | 35 | M | 182 | 71 | 21.4 |  |  |
|  | 32 | M | 176 | 67 | 21.6 |  |  |
|  | 32 | F | 170 | 64 | 22.1 |  |  |
|  | 22 | M | 190 | 70 | 19.4 |  |  |
|  | 38 | M | 170 | 68 | 23.5 |  |  |
|  | 34 | F | 160 | 55 | 21.5 |  |  |
| Older untrained | 59 | M | 164 | 53 | 19.7 | Osteoporosis | Perindopril, Amlodipine, Etidronate, Adcal |
|  | 91 | F |  |  |  | Osteoporosis, Diverticulitis - Hartmann's bowel operation | Furosemide, Paracetamol, Colecalciferol, Docusate |
|  | 77 | M |  |  |  | Osteoporosis | Paracetamol, Tramadol, omeprazol, renetadine, mirapixin, procloperizone |
|  | 82 | F | 152 | 57 | 24.7 | Osteoporosis, Hypertension, Rheumatoid arthritis, type 2 diabetes | Metformin, denusomab, levothyroxine, allopurinol |
|  | 88 | F | 156 | 64 | 26.3 | Osteoporosis, Breast cancer, left mastectomy in remission | Levothroxine, atenolol |
|  | 77 | F |  | 61 |  | Osteoporosis, Rheumatoid arthritis | Sulfasalazine |
| Master cyclists (older trained) | 71 | M | 180 | 83 | 25.7 |  |  |
|  | 76 | F | 157 | 58 | 23.4 |  |  |
|  | 79 | F | 159 | 53 | 21.1 |  |  |
|  | 75 | M | 172 | 62 | 20.8 |  |  |
|  | 73 | M | 170 | 70 | 24.0 |  |  |
|  | 79 | F |  |  |  |  |  |

The laborious and time-consuming nature of recruiting human participants and acquiring their samples for longitudinal studies makes cross-sectional analyses more feasible. To confirm our findings, a longitudinal study of myonuclear structure and function in samples serially acquired from active and inactive individuals throughout their lifespan would be required. This would provide a comprehensive account of the effects of active and inactive ageing on skeletal muscle myonuclei, and a causal relationship between changes in myonuclear parameters and muscle function.

In summary, our data suggest that exercise is associated with profound alterations in nuclear structure and mechanics in human primary muscle fibres regardless of age. In line with this, exercise resulted in increased Lamin A expression and myonuclear stiffness in mice. Future investigations into the potential role of myonuclear mechanotransduction in exercise and ageing would further our understanding of skeletal muscle physiology and offer new insights into improving human healthspan.

## Methods

### Participant characteristics and ethics

Four mixed gender groups were recruited to participate in the current study (n = 6 per group). These groups were: younger untrained healthy (YU) (33 ± 9.5 years), younger trained marathon runners (YT) (32 ± 5.4 years), older untrained individuals (OU) (79 ± 11.3 years), and older highly trained cyclists (OT) (75.5 ± 3.2 years) (Table 1). The YU group was considered healthy, but not necessarily sedentary, as two of the participants had been participating in low-level recreational sport activities (<2 sessions/week) at the time of the study. Thus, the young cohort consisted of low-level physically active and sedentary individuals. Participants were considered healthy if they met the criteria outlined by Greig et al. (Greig *et al*., 1994). The exclusion criteria from a healthy classification were smoking or consuming alcohol excessively, known hypertension or other cardiovascular, musculoskeletal, or neurological conditions, or if they were on any medication (acute or chronic). The YT group consisted of trained marathon runners (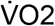 peak 56.7 ± 6.6 ml.kg.min^-1^, mean ± SD). In the YT group, the fastest running times (mean ± SD) in the previous 18 months over marathon, half marathon and 5 km distances were 204.5 ± 14.2 min, 88.5 ± 3.3 min, and 19.8 ± 1.3 min, respectively. The OU group, used as a model for muscle disuse in old age, was a previously characterised cohort, who underwent dynamic hip screw insertion surgery. The patients completed a basic physical health questionnaire and were considered eligible if they did not suffer from neuromuscular disease, although some had underlying health conditions (Table 1). The OT group consisted of previously characterised individuals (Pollock *et al*., 2015) who were amateur master endurance cyclists. Master cyclists were included if they were able to cycle 100 km in under 6.5 hours (males) or 60 km in under 5.5 hours (females). Participants must have had completed this distance within the specified time on two occasions in the three weeks prior to the date of participation in the study.

### Obtaining and processing skeletal muscle samples and isolating single muscle fibres

Vastus lateralis samples were obtained as previously described (Pollock *et al*., 2018). Approximately 60mg of the biopsy sample was then placed in relaxing solution (77.63 mM KCL, 10 mM imidazole, 2 mM MgCl_2_, 2 mM EGTA, 4.05 mM ATP in distilled water, pH 7.0) in a petri dish on ice. Following excision, muscle samples (submerged in relaxing solution in a petri dish) were divided into bundles of approximately 100 muscle fibres using forceps under a stereo microscope (Zeiss, Stemi 2000-C) with a separate light source (Zeiss Stereo CL 1500 ECO). The ends of the bundles were then tied onto glass capillary tubes using surgical silk (LOOK SP102) and stretched to approximately 110% of the original length. These bundles were subsequently placed into 1.5 ml Eppendorf tubes, containing skinning solution (relaxing solution with 50% (v/v) glycerol), at 4°C for 48 h to permeabilise the muscle fibres by disrupting the lipid bilayer of the sarcolemma, leaving myofilaments, intermediate filaments, and nuclear envelope intact (Konigsberg *et al*., 1975; Wood *et al*., 1975; Frontera & Larsson, 1997; Stienen, 2000). Samples were then treated in ascending gradients of sucrose dissolved in relaxing solution (0.5 M, 1 M, 1.5 M, 2 M) for 30 minutes to prevent cryodamage (Frontera & Larsson, 1997). In a petri dish containing 2 M sucrose, fibres were then removed from the glass capillary tubes before being placed in cryovials and snap-frozen in liquid nitrogen.

For immunofluorescence and nuclear mechanics experiments, muscle fibre bundles were placed in descending concentrations of sucrose dissolved in relaxing solution, for 30 minutes in each solution (2 M, 1.5 M, 1 M, 0.5 M, 0 M). Samples were transferred to skinning solution at −20°C until the day of an experiment. To isolate single muscle fibres, muscle bundles were placed in skinning solution in a petri dish on an ice block. One end of the muscle bundle was held using extra fine forceps, whilst single fibres were pulled from the end of the bundle. During this process, care was taken to restrict contact to the ends of fibres as much as possible to avoid damage. To normalise muscle fibre tension and orientation, muscle fibres were mounted on half-split grid for transmission electron microscopy (TEM) glued to a coverslip (Ross *et al*.,2017; Levy *et al*., 2018). Fibres were then immunostained and imaged or analysed by a nanoindenter to assess nuclear mechanics.

### Immunostaining, imaging, and analysis of single muscle fibres

The first steps of each staining protocol were fixing in 4% PFA for 15 min and permeabilising in 0.2% triton for 10 min. When primary antibodies were used, fibres were blocked using 10% goat serum (Sigma-Aldrich, G9023) in PBS for 1 hour at room temperature before incubation in primary antibody solution overnight at 4°C. Muscle fibres were then incubated in a solution containing direct stains and secondary antibodies for 1 h (see list of primary and secondary in Table 2). Finally, fibres were mounted in Fluoromount-G^®^or DAKO mounting medium. Between each step of staining protocols, fibres were washed four times in PBS.

**Table 2.**
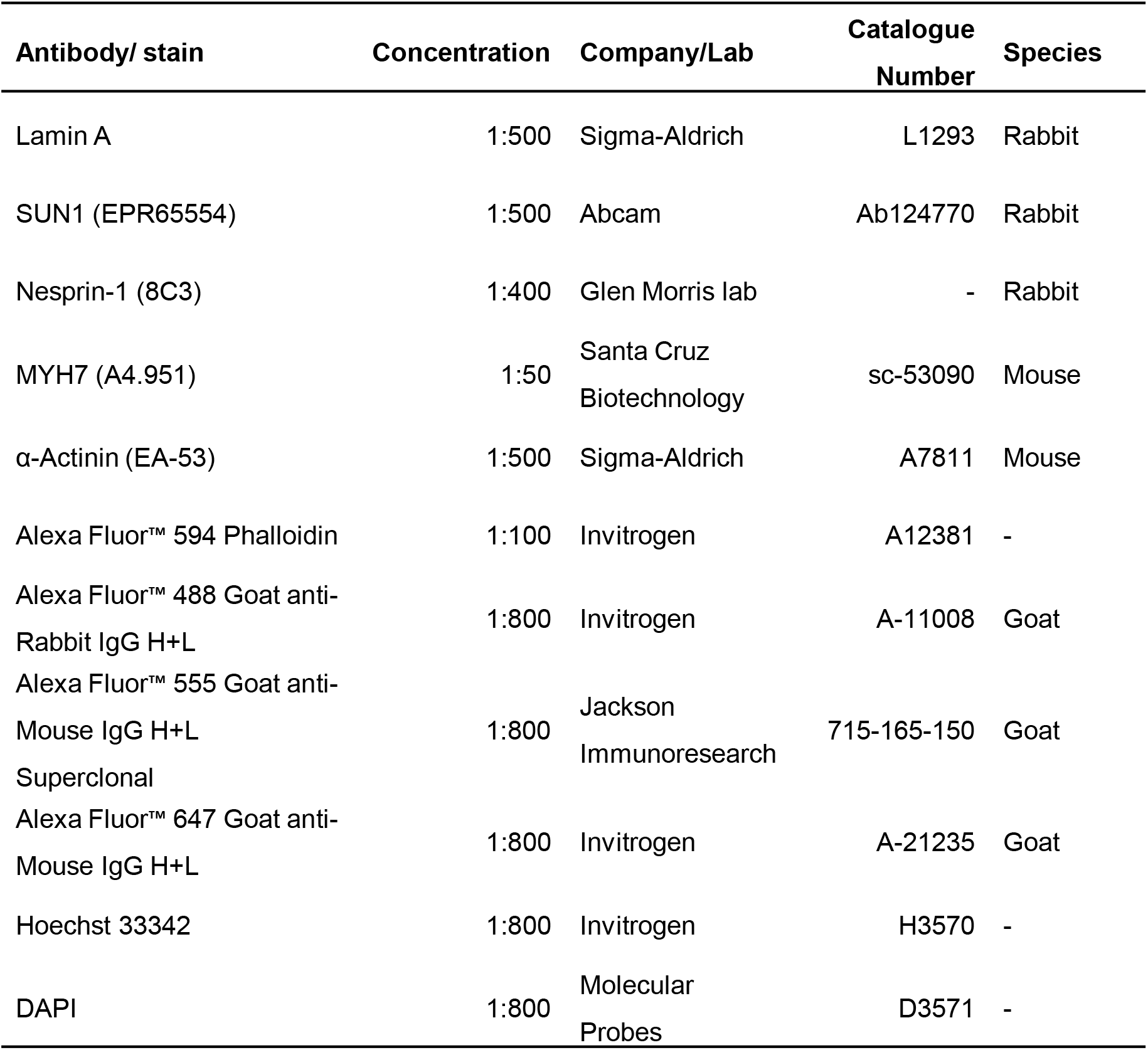
Antibodies and direct stains used in immunofluorescence experiments.

For two-dimensional analysis of myonuclear shape, single plane or z-stack images (1 μm Z increments) were acquired using a 40x air objective and a Zeiss Axiovert 200 microscope system. Two-dimensional myonuclear shape parameters (nuclear area and aspect ratio) were quantified using Fiji software, as previously described (Schindelin *et al*., 2012; Battey *et al*.,2022). Single plane or maximum intensity projection images were processed with a rolling ball background subtraction (150 pixels), gaussian blur filter (2 pixels radius) and despeckle function before thresholding the DAPI signal (initially with ‘Otsu dark’ setting then adjusting as necessary). Laterally located nuclei (i.e. positioned around the sides of muscle fibres) were excluded from analysis as nuclei in this position are orientated perpendicular, rather than facing the objective lens.

For three-dimensional analysis of myonuclear shape and Lamin A organisation, Z-stack images (0.2 μm Z increments) were acquired using a 60x oil objective and a Nikon spinning disk confocal microscope system. To quantify three-dimensional shape parameters (sphericity; skeletal length/diameter, referred to as 3D aspect ratio), the DAPI signal was thresholded and analysed using Volocity software (Perkin Elmer). Representative images were produced by generating standard deviation pixel projections of Z-stacks in Fiji.

To visualise and analyse Lamin A at super-resolution level, Z-stack images (0.1 μm increments) were acquired with a 100x oil objective (numerical aperture 1.5) and a Nikon instant Structured Illumination Microscope (iSIM) system. At least six fibres were imaged per individual, with each image including 1-7 myonuclei. To improve contrast and resolution (by two-fold compared to confocal microscopy), iSIM images were deconvolved using inbuilt algorithms in Nikon Elements software (3D Blind algorithm with 15 iterations and spherical aberration correction) (York *et al*., 2013; Curd *et al*., 2015). The organisation of the nuclear lamina was analysed using Fiji software. Line scan analysis of Lamin A staining was performed by using the plot profile tool. Nuclear lamina deposition (arbitrary units) was quantified by using a full width at half maximum macro to fit a Gaussian curve to pixel intensity profiles of Lamin A stains. Measurements using the tool were taken in mid-focal planes, with an average taken from a minimum of 6 measurements per nuclei. For nuclear invagination length analysis, a Fiji plugin called Ridge Detection was used (Steger, 1998).

### Assessment of myonuclear mechanics in single muscle fibres

To assess myonuclear mechanics in single muscle fibres, the extent of nuclear deformation with increasing fibre tension was quantified (Shah & Lieber, 2003; Chapman *et al*., 2014). Images were acquired as before for 2D analysis, and sarcomere length was correlated with nuclear aspect ratio, to assess myonuclear shape at various extents of muscle fibre stretch.

To determine stiffness, elastic, and viscosity properties of myonuclei, nanoindentation was carried out on mounted single muscle fibres. Experiments were performed using an Optics11 Chiaro nanoindenter attached to a Leica DMI-8 inverted microscope. Mounted muscle fibres were stained with Hoescht to locate myonuclei, and only nuclei at the nearest surface of the muscle fibre to the indenter were analysed. Nanoindentation was performed with a 9 μm diameter spherical probe (2.1 μm contact radius at 0.5 μm indentation depth, corresponding to approximately half the nuclear radius, whereby an indentation at 0.5 μm depth measures primarily the nuclear vs the cytoskeletal contribution to the stiffness (Guerrero *et al*., 2019)).

Approach and retraction speeds were set to 500 nm/s. The Hertzian contact model was used to fit the load-indentation data for calculation of Young’s modulus.

Dynamic Mechanical Analysis (DMA), which uses a cyclic motion with frequency while controlling displacement or load, was used to calculate the frequency-dependent storage modulus (*E’*), loss modulus (*E”*), and the dissipation factor tan delta (*E’/E”*) of myonuclei, corresponding to the elastic properties of nuclei. 1, 2, 4 and 10 Hz frequencies were used (Table 3).

**Table 3.** Dynamic mechanical analysis parameters for nanoindentation.

| Frequency (Hz) | Indentation depth (nm) | Periods | Time (s) |
| --- | --- | --- | --- |
| 1 | 500 | 5 | 2 |
| 2 | 500 | 5 | 2 |
| 4 | 500 | 20 | 2 |
| 10 | 500 | 20 | 2 |

## Mouse high intensity interval training programme

12 C57BL/6 mice were trained on treadmill over 8 weeks, with a complementary sedentary group left in cages for an equivalent time-period. At the end of the 8 weeks, *tibialis anterior* muscle was excised from both legs of each mouse. Muscle from one leg was placed in skinning solution before cryopreservation and storage at −80 °C for later analysis through nanoindentation. Muscle from the contralateral leg was snap frozen in liquid nitrogen for western blot analysis.

### Treadmill familiarisation, determination of peak running velocity and training programme

Mice were familiarised to the treadmill with five days of low intensity running (5 cm/s on the first day, increasing the speed by 5 cm/s on the second, third, and fifth day, to end with a speed of 20 cm/s). Treadmill incline was set at 0° on day one, 5° on days two and three, and 10° on day five. Peak running velocity (VPeak) was determined to estimate maximal aerobic capacity and allow standardisation of the intensity of running during the training programme. Exercise prescription based on VPeak and 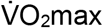 result in similar aerobic adaptations in both humans and mice (Manoel *et al*., 2017; Picoli *et al*., 2018). VPeak was determined based on the method outlined by Picoli et al. (Picoli *et al*., 2018), adapted to incorporate a ramp, rather than incremental, increase in running speed, as suggested by Ayachi et al. (Ayachi *et al*., 2016). Testing commenced with a warm-up for 4 min at 10 cm/s, before increasing the speed gradually to 19 cm/s over the next minute (approximately 1 cm/s every 6.5 seconds). Running speed was then increased by 1 cm/s every 20 seconds until exhaustion, characterised by incapacity to keep running for more than 5 s (Mille-Hamard *et al*., 2012).

Mice ran four times per week for eight weeks, based on a programme that showed ~50% improvement in 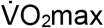 (Kemi *et al*., 2002; Høydal *et al*., 2007). After a warm-up at 10 cm/s for 6 min, mice ran at approximately 80-90% Vpeak for three bouts of 8 min intermitted by 2 min active recovery at 50-60% Vpeak. Table 4 shows the speed and inclines of the training programme. Muscle was excised from mice (sacrificed by cervical dislocation) 72 hours after the final exercise session to exclude potential confounding effects observed acutely after exercise (Carmichael *et al*., 2005; Neubauer *et al*., 2014).

**Table 4.**
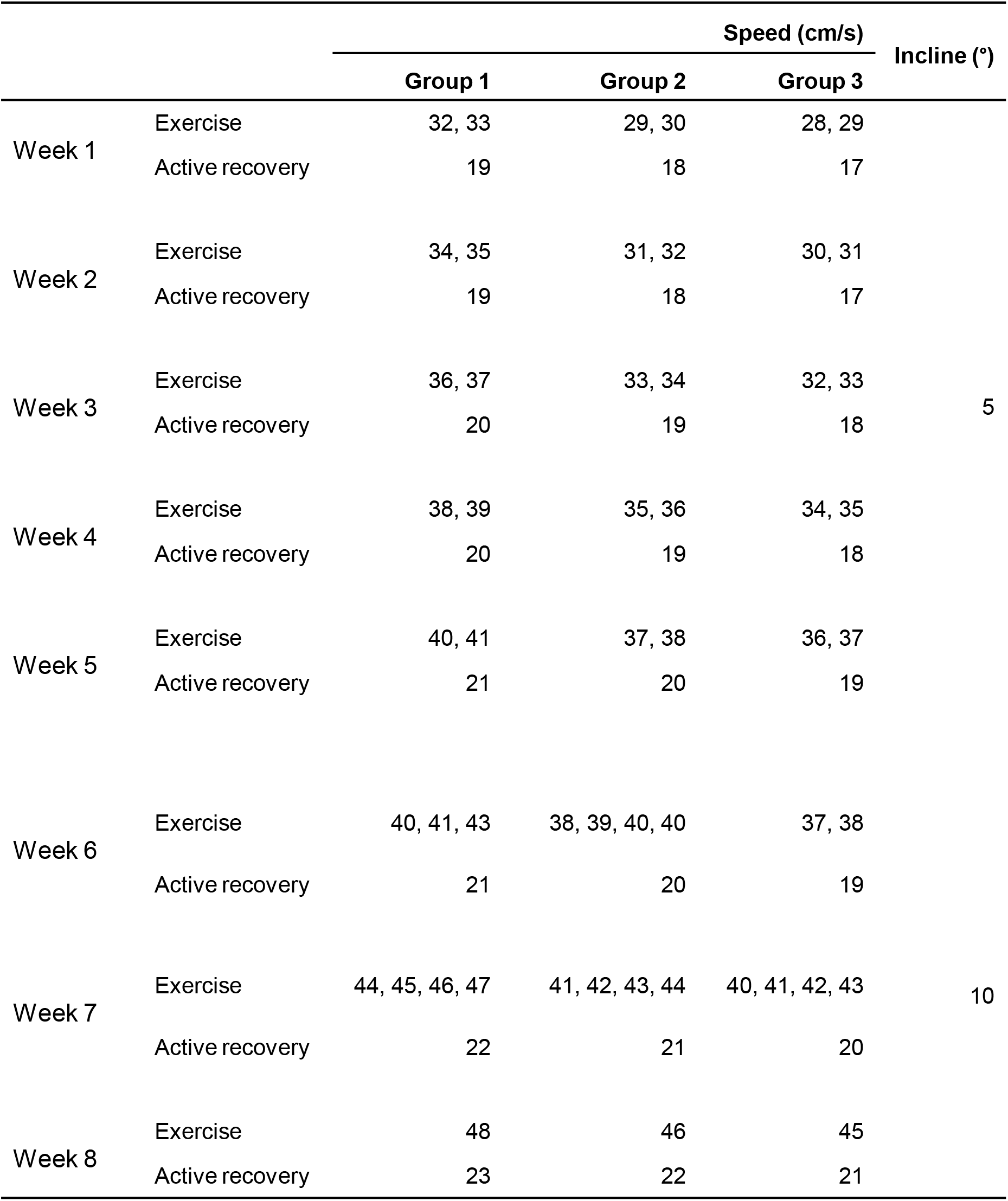
Speed and incline of treadmill throughout mouse high intensity interval training programme.

### Quantification of mouse skeletal muscle protein content by western blotting

Tissue samples were lysed in 5M Urea, 2M Thiourea, 3% SDS, 75mM DTT, 0.03% Bromophenol Blue, and 0.05M Tris HCl, and homogenised in a Precellys 24 tissue homogeniser machine kept at 4°C. Samples were then sonicated for additional homogenisation and shearing of DNA. For western blotting, frozen tissue lysates were thawed and proteins linearised in a heating block at 95°C for 8 minutes, before loading on 4-12% Bis-Tris gels. Proteins were transferred onto nitrocellulose membranes and blocked in 5% non-fat milk powder in Tris Buffered Saline with 0.1% Tween20 (TBS-T) at 4°C for 1 hour. The membranes were then incubated with primary antibodies overnight at 4°C, washed in TBS-T 4 times, and subsequently incubated with secondary antibodies for 1 hour. After 4 washes, membranes imaged using a LI-COR Odyssey^®^ CLx imaging system. Antibodies used for western blotting are outlined in Table 5.

**Table 5.** Antibodies used for western blotting.

| **Antibody** | **Company/Lab** | **Catalogue Number** | **Species** |
| --- | --- | --- | --- |
| Nesprin-1 (8C3) | Glen Morris Lab | - | Mouse |
| Nesprin-2 | Didier Hodzic Lab | - | Rabbit |
| SUN1 | Abcam | ab103021 | Rabbit |
| SUN2 | Didier Hodzic Lab | - | Rabbit |
| Lamin A/C | Larry Gerace Lab | - | Rabbit |
| Lamin B1 | Larry Gerace Lab | - | Rabbit |
| Lamin B2 | Abcam | ab151735 | Rabbit |
| LAP1 | William Dauer Lab | - | Rabbit |
| Lem2 | Atlas Antibodies | HPA017340 | Rabbit |
| Emerin | Santa Cruz<br>Biotechnologies | FL-254 | Rabbit |
| HP1α | Cell Signaling<br>Technology | 2616S | Rabbit |
| HP1γ | Cell Signaling<br>Technology | 2619S | Rabbit |
| Histone H3K9me2 | Active Motif | 39041 | Rabbit |
| Histone H3K9me3 | Novus Biologicals | NBP1-30141 | Mouse |
| γH2AX | Cell Signaling<br>Technology | 9718S | Rabbit |
| Histone H3 (D1H2) | Cell Signaling<br>Technology | 4499T | Rabbit |
| GAPDH (D4C6R) | Cell Signaling<br>Technology | 97166S | Mouse |
| IRDye® 680RD Donkey<br>anti-Mouse IgG<br>secondary antibody | LI-COR Biosciences | 926-68072 | Donkey |
| IRDye® 800CW Donkey<br>anti-Rabbit IgG<br>secondary antibody | LI-COR Biosciences | 926-32213 | Donkey |

## Statistics

Based on an expected mean difference of 15% between young and elderly individuals, an effect size of 1.8, α = 0.05 and power (1-β) = 0.8, the required sample size was determined as six individuals per group. This studied was powered based on nuclear aspect ratio as the primary end-point measurement. Because there were no studies on nuclear shape changes in human muscle fibres, the study was powered based on data showing increased aspect ratio of this magnitude in muscle fibres from mice expressing mutant lamins compared to control mice (Earle *et al*., 2020). To analyse whether an overall significant difference was present between muscle fibres from the different human groups, two-way analysis of variance (ANOVA) tests were carried out, followed by post-hoc Tukey tests to specify between which myonuclei or muscle fibres group differences existed. Mean values for each individual were used for two-way ANOVA. For correlation analyses, a simple linear regression was performed to test if slopes were significantly non-zero and nonlinear straight-line regression analyses were performed to compare the slopes of different conditions. Two tailed t-tests were carried out to analyse differences in protein concentrations, and Young’s modulus between myonuclei from exercise trained and untrained mice. Mean values for each mouse were used for t-tests. With categorical data, individual measurements (myonuclei or muscle fibres) and mean values calculated from these measurements were plotted in the same graph, using SuperPlots(Lord *et al*., 2020). Individual measurements were plotted as smaller grey points and overall means of each individual or mouse are plotted as larger coloured points. For all statistical tests, p < 0.05 indicated significance and p < 0.07 was taken to indicate a trend (* P < 0.05, ** P < 0.01, *** P < 0.001). Data are presented as mean ± SD except mean ± SEM in Figure 5. All data were statistically analysed using Prism 9 (GraphPad).

## Precision and reproducibility of methods

The same image processing and initial thresholding parameters were used for each image for quantification of nuclear shape, nuclear organisation, lamina deposition, nuclear envelope protein organisation. To quantify the precision and accuracy of the methods used, repeat measurements were taken and the standard deviation and coefficient of variation were calculated. Three repeat measurements of three nuclei were carried out for nuclear aspect ratio and nuclear area. The mean of the standard deviation and coefficient of variation values were then calculated to give a single standard deviation and coefficient of variation value for nuclear aspect ratio and nuclear area. Five repeat measurements of sarcomere length were taken, and five repeat full width at half-maximum measurements of the nuclear lamina were carried out for assessment of the precision of lamina deposition. The standard deviation and coefficient of variation values were 0.02 and 0.67 for nuclear aspect ratio, 1.19 and 1.46 for nuclear area, 0.07 and 0.28 for sarcomere length, and 0.004 and 1.53 for lamina deposition, respectively.

## Study approval

Prior to participation, written informed consent was obtained from all subjects. Procedures were approved by the Fulham Research Ethics Committee in London (12/LO/0457), Westminster Ethics Committee in London (12/LO/0457) or Liverpool John Moores ethics committee (H17SPS012) and conformed to the Declaration of Helsinki. All human tissues were collected, stored, and analysed in accordance with the Human Tissue Act. Procedures for the mouse study were performed in accordance with the Guidance on the Operation of the Animals (Scientific Procedures) Act, 1986 (UK Home Office).

## Author Contributions

MJS and JO contributed to the conception of the work. EB, JAR, MJS and JO designed experiments. EB, JAR, AH, DGSW, YH, TI, JO and MJS did the acquisition, analysis, and interpretation of data. RDP, MK, JNP, GLC, NRL, SDRH and JO recruited human participants and collected human muscle biopsy samples. EB and MJS wrote the first draft of the manuscript. EB, JAR, AH, RDP, MK, GME, NRL, TI, SDRH, JO and MJS contributed to the manuscript and revised it critically. EB, JAR, AH, DGSW, YL, RDP, MK, JNP, GLC, GME, NRL, TI, SDRH, JO and MJS approved the final version of the manuscript to be published. EB, JAR, AH, DGSW, YL, RDP, MK, JNP, GLC, GME, NRL, TI, SDRH, JO and MJS agreed on all aspects of the work.

## Data availability statement

Individual datapoints (*n* ≤ 30) are included in the figures. Data from the study will be made available upon reasonable request.

## Acknowledgements

Julien Ochala and Edmund Battey are funded by the Medical Research Council of the UK (MR/S023593/1). Matthew J. Stroud is supported by British Heart Foundation Intermediate Fellowship: FS/17/57/32934 and King’s BHF Centre for Excellence Award: RE/18/2/34213. Stephen D. R. Harridge, Norman R. Lazarus and Ross D. Pollock were funded by the Bupa Foundation. Michaeljohn Kalakoutis was funded by a King’s College London PhD studentship.

## Figure legends

**Supplementary Figure 1:**
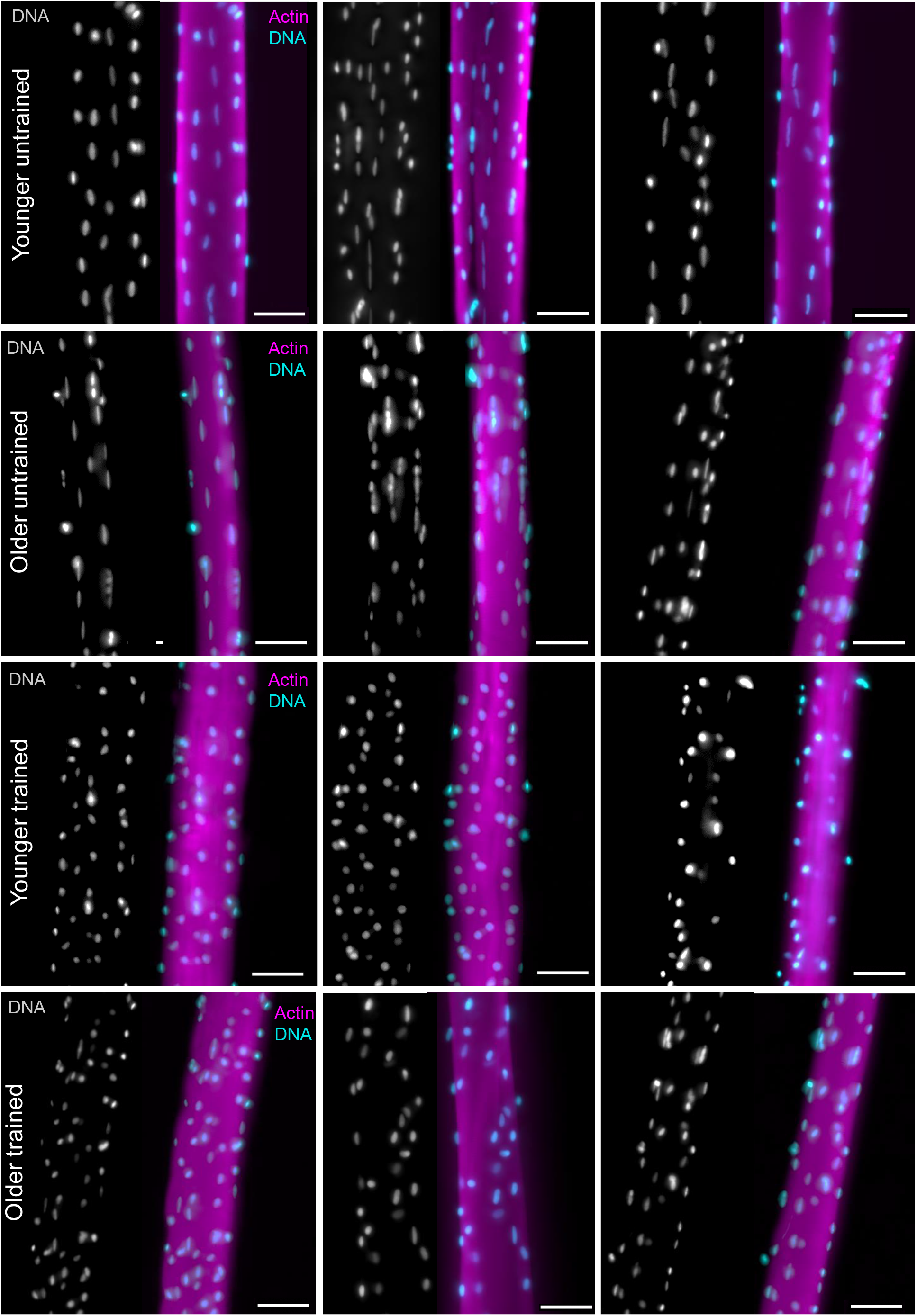
Representative images of myonuclear organisation in younger and older untrained and trained individuals. Representative images of vastus lateralis muscle fibres isolated from younger untrained, older untrained, younger trained and older trained stained with DAPI (cyan) and Phalloidin (magenta) to visualise myonuclei and Actin, respectively. Scale bars, 50 μm.

## Notes

**Conflict-of-interest statement:**the authors have declared that no conflict of interest exists.

### Competing Interest Statement

The authors have declared no competing interest.

### Summary of Updates

Figures have been integrated into the text. Yu Han added as a co-author and the author contributions have been updated accordingly.

## References

United Nations (2022). World Population Prospects 2022 Summary of Results.

Attwaters M & Hughes SM (2022). Cellular and molecular pathways controlling muscle size in response to exercise. FEBS Journal 289, 1428–1456.

Ayachi M, Niel R, Momken I, Billat VL & Mille-Hamard L (2016). Validation of a ramp running protocol for determination of the true VO2max in mice. Front Physiol; DOI: 10.3389/FPHYS.2016.00372/FULL.

Banerjee I, Zhang J, Moore-Morris T, Pfeiffer E, Buchholz KS, Liu A, Ouyang K, Stroud MJ, Gerace L, Evans SM, McCulloch A & Chen J (2014). Targeted Ablation of Nesprin 1 and Nesprin 2 from Murine Myocardium Results in Cardiomyopathy, Altered Nuclear Morphology and Inhibition of the Biomechanical Gene Response. PLoS Genet 10, e1004114.

Battey E, Furrer R, Ross J, Handschin C, Ochala J & Stroud MJ (2022). PGC-1α regulates myonuclear accretion after moderate endurance training. J Cell Physiol 237, 696–705.

Battey E, Stroud MJ & Ochala J (2020). Using nuclear envelope mutations to explore age-related skeletal muscle weakness. Clin Sci 134, 2177–2187. Available at: https://portlandpress.com/clinsci/article-abstract/134/16/2177/226196 [Accessed February 16, 2021].

Bonne G, Schwartz K, Barletta MR di, Varnous S, Bécane H-M, Hammouda E-H, Merlini L, Muntoni F, Greenberg CR, Gary F, Urtizberea J-A, Duboc D, Fardeau M & Toniolo D (1999). Mutations in the gene encoding lamin A/C cause autosomal dominant Emery-Dreifuss muscular dystrophy. Nat Genet 21, 285–288.

Brown GC (2015). Living too long. EMBO Rep 16, 137–141.

Buxboim A, Swift J, Irianto J, Spinler KR, Dingal PCDP, Athirasala A, Kao YRC, Cho S, Harada T, Shin JW & Discher DE (2014). Matrix elasticity regulates lamin-A,C phosphorylation and turnover with feedback to actomyosin. Curr Biol 24, 1909–1917.

Carmichael MD, Davis JM, Murphy EA, Brown AS, Carson JA, Mayer E & Ghaffar A (2005). Recovery of running performance following muscle-damaging exercise: Relationship to brain IL-1β. Brain Behav Immun 19, 445–452.

Chapman MA, Zhang J, Banerjee I, Guo LT, Zhang Z, Shelton GD, Ouyang K, Lieber RL & Chen J (2014). Disruption of both nesprin 1 and desmin results in nuclear anchorage defects and fibrosis in skeletal muscle. Hum Mol Genet 23, 5879–5892.

Cho S, Irianto J & Discher DE (2017). Mechanosensing by the nucleus: From pathways to scaling relationships. Journal of Cell Biology 216, 305–315.

Crisp M, Liu Q, Roux K, Rattner JB, Shanahan C, Burke B, Stahl PD & Hodzic D (2006). Coupling of the nucleus and cytoplasm: Role of the LINC complex. Journal of Cell Biology 172, 41–53.

Cupesi M, Yoshioka J, Gannon J, Kudinova A, Stewart CL & Lammerding J (2010). Attenuated hypertrophic response to pressure overload in a lamin A/C haploinsufficiency mouse. J Mol Cell Cardiol 48, 1290–1297.

Curd A, Cleasby A, Makowska K, York A, Shroff H & Peckham M (2015). Construction of an instant structured illumination microscope. Methods 88, 37–47.

Earle AJ, Kirby TJ, Fedorchak GR, Isermann P, Patel J, Iruvanti S, Moore SA, Bonne G, Wallrath LL & Lammerding J (2020). Mutant lamins cause nuclear envelope rupture and DNA damage in skeletal muscle cells. Nat Mater 19, 464–473.

Fischer M, Rikeit P, Knaus P & Coirault C (2016). YAP-mediated mechanotransduction in skeletal muscle. Front Physiol 7, 41.

Frontera WR & Larsson L (1997). Contractile studies of single human skeletal muscle fibers: A comparison of different muscles, permeabilization procedures, and storage techniques. Muscle Nerve 20, 948–952.

Frost B (2016). Alzheimer’s disease: An acquired neurodegenerative laminopathy. Nucleus 7, 275–283.

Gerhart-Hines Z, Rodgers JT, Bare O, Lerin C, Kim SH, Mostoslavsky R, Alt FW, Wu Z & Puigserver P (2007). Metabolic control of muscle mitochondrial function and fatty acid oxidation through SIRT1/PGC-1α. EMBO J 26, 1913–1923.

Goldman RD, Shumaker DK, Erdos MR, Eriksson M, Goldman AE, Gordon LB, Gruenbaum Y, Khuon S, Mendez M, Varga R & Collins FS (2004). Accumulation of mutant lamin A causes progressive changes in nuclear architecture in Hutchinson–Gilford progeria syndrome. Proceedings of the National Academy of Sciences 101, 8963–8968.

Greig CA, Young A, Skelton DA, Pippet E, Butler FMM & Mahmud SM (1994). Exercise studies with elderly volunteers. Age Ageing 23, 185–189.

Guerrero CR, Garcia PD & Garcia R (2019). Subsurface Imaging of Cell Organelles by Force Microscopy. ACS Nano 13, 9629–9637.

Guilluy C, Osborne LD, van Landeghem L, Sharek L, Superfine R, Garcia-Mata R & Burridge K (2014). Isolated nuclei adapt to force and reveal a mechanotransduction pathway in the nucleus. Nat Cell Biol 16, 376–381.

Gurd BJ (2011). Deacetylation of PGC-1a by SIRT1: Importance for skeletal muscle function and exercise-induced mitochondrial biogenesis. Applied Physiology, Nutrition and Metabolism 36, 589–597. Available at: www.nrcresearchpress.com [Accessed April 17, 2022].

Guthold R, Stevens GA, Riley LM & Bull FC (2018). Worldwide trends in insufficient physical activity from 2001 to 2016: a pooled analysis of 358 population-based surveys with 1·9 million participants. Lancet Glob Health 6, e1077–e1086.

Ho CY, Jaalouk DE, Vartiainen MK & Lammerding J (2013). Lamin A/C and emerin regulate MKL1-SRF activity by modulating actin dynamics. Nature 497, 507–511.

Hornberger TA & Esser KA (2004). Mechanotransduction and the regulation of protein synthesis in skeletal muscle. Proceedings of the Nutrition Society 63, 331–335.

Høydal MA, Wisløff U, Kemi OJ & Ellingsen Ø (2007). Running speed and maximal oxygen uptake in rats and mice: practical implications for exercise training. European journal of cardiovascular prevention and rehabilitation 14, 753–760.

Janin A, Bauer D, Ratti F, Millat G & Méjat A (2017). Nuclear envelopathies: A complex LINC between nuclear envelope and pathology. Orphanet J Rare Dis 12, 1–16. Available at: https://ojrd.biomedcentral.com/articles/10.1186/s13023-017-0698-x [Accessed April 18, 2022].

Jones SW, Hill RJ, Krasney PA, O’Conner B, Peirce N & Greenhaff PL (2004). Disuse atrophy and exercise rehabilitation in humans profoundly affects the expression of genes associated with the regulation of skeletal muscle mass. The FASEB Journal 18, 1025–1027.

Kalukula Y, Stephens AD, Lammerding J & Gabriele S (2022). Mechanics and functional consequences of nuclear deformations. Nature Reviews Molecular Cell Biology 20221–20.

Kemi OJ, Loennechen JP, Wisløff U & Ellingsen Y (2002). Intensity-controlled treadmill running in mice: Cardiac and skeletal muscle hypertrophy. J Appl Physiol 93, 1301–1309.

Kirby TJ (2019). Mechanosensitive pathways controlling translation regulatory processes in skeletal muscle and implications for adaptation. J Appl Physiol 127, 608–618.

Kirby TJ & Lammerding J (2018). Emerging views of the nucleus as a cellular mechanosensor. Nat Cell Biol 20, 373–381.

Konigsberg UR, Lipton BH & Konigsberg IR (1975). The regenerative response of single mature muscle fibers isolated in vitro. Dev Biol 45, 260–275.

Lammerding J, Fong LG, Ji JY, Reue K, Stewart CL, Young SG & Lee RT (2006). Lamins a and C but not lamin B1 regulate nuclear mechanics. Journal of Biological Chemistry 281, 25768–25780.

Lazarus NR & Harridge SDR (2017). Declining performance of master athletes: silhouettes of the trajectory of healthy human ageing? Journal of Physiology 595, 2941–2948.

Lazarus NR, Lord JM & Harridge SDR (2019). The relationships and interactions between age, exercise and physiological function. Journal of Physiology 597, 1299–1309.

Levy Y, Ross JA, Niglas M, Snetkov VA, Lynham S, Liao C-Y, Puckelwartz MJ, Hsu Y-M, McNally EM, Alsheimer M, Harridge SDR, Young SG, Fong LG, Español Y, Lopez-Otin C, Kennedy BK, Lowe DA & Ochala J (2018). Prelamin A causes aberrant myonuclear arrangement and results in muscle fiber weakness. JCI Insight 3, 1–18.

Liang Y, Chiu PH, Yip KY & Chan SY (2011). Subcellular localization of SUN2 is regulated by lamin a and Rab5. PLoS One; DOI: 10.1371/journal.pone.0020507.

Little JP, Safdar A, Wilkin GP, Tarnopolsky MA & Gibala MJ (2010). A practical model of low-volume high-intensity interval training induces mitochondrial biogenesis in human skeletal muscle: potential mechanisms. J Physiol 588, 1011–1022.

Lord SJ, Velle KB, Dyche Mullins R & Fritz-Laylin LK (2020). SuperPlots: Communicating reproducibility and variability in cell biology. Journal of Cell Biology; DOI: 10.1083/JCB.202001064.

Maggi L, Carboni N & Bernasconi P (2016). Skeletal Muscle Laminopathies: A Review of Clinical and Molecular Features. Cells 5, 33.

Manoel F, da Silva D, Lima J & Machado F (2017). Peak velocity and its time limit are as good as the velocity associated with VO2max for training prescription in runners. Sports Med Int Open 01, E8–E15.

Maurer M & Lammerding J (2019). The Driving Force: Nuclear Mechanotransduction in Cellular Function, Fate, and Disease. Annu Rev Biomed Eng 21, annurev-bioeng-060418-052139.

McClintock D, Gordon LB & Djabali K (2006). Hutchinson-Gilford progeria mutant lamin A primarily targets human vascular cells as detected by an anti-Lamin A G608G antibody. Proc Natl Acad Sci U S A 103, 2154–2159.

Merideth MA et al. (2008). Phenotype and course of Hutchinson-Gilford progeria syndrome. New England Journal of Medicine 358, 592–604.

Mille-Hamard L, Billat VL, Henry E, Bonnamy B, Joly F, Benech P & Barrey E (2012). Skeletal muscle alterations and exercise performance decrease in erythropoietin-deficient mice: A comparative study. BMC Med Genomics; DOI: 10.1186/1755-8794-5-29.

Murach KA, Mobley CB, Zdunek CJ, Frick KK, Jones SR, McCarthy JJ, Peterson CA & Dungan CM (2020). Muscle memory: myonuclear accretion, maintenance, morphology, and miRNA levels with training and detraining in adult mice. J Cachexia Sarcopenia Muscle 11, 1705–1722.

Neubauer O, Sabapathy S, Ashton KJ, Desbrow B, Peake JM, Lazarus R, Wessner B, Cameron-Smith D, Wagner KH, Haseler LJ & Bulmer AC (2014). Time course-dependent changes in the transcriptome of human skeletal muscle during recovery from endurance exercise: From inflammation to adaptive remodeling. J Appl Physiol 116, 274–287.

Nikitara K, Odani S, Demenagas N, Rachiotis G, Symvoulakis E & Vardavas C (2021). Prevalence and correlates of physical inactivity in adults across 28 European countries. Eur J Public Health 31, 840–845.

Osmanagic-Myers S, Dechat T & Foisner R (2015). Lamins at the crossroads of mechanosignaling. Genes Dev 29, 225–237.

Owens DJ, Messéant J, Moog S, Viggars M, Ferry A, Mamchaoui K, Lacène E, Roméro N, Brull A, Bonne G, Butler-Browne G & Coirault C (2021). Lamin-related congenital muscular dystrophy alters mechanical signaling and skeletal muscle growth. Int J Mol Sci 22, 1–22.

Park YE, Hayashi YK, Goto K, Komaki H, Hayashi Y, Inuzuka T, Noguchi S, Nonaka I & Nishino I (2009). Nuclear changes in skeletal muscle extend to satellite cells in autosomal dominant Emery-Dreifuss muscular dystrophy/limb-girdle muscular dystrophy 1B. Neuromuscular Disorders 19, 29–36.

Piccus R & Brayson D (2020). The nuclear envelope: LINCing tissue mechanics to genome regulation in cardiac and skeletal muscle. Biol Lett; DOI: 10.1098/rsbl.2020.0302. Available at: https://royalsocietypublishing.org/doi/10.1098/rsbl.2020.0302 [Accessed September 20, 2022].

Picoli C de C, Romero PV da S, Gilio GR, Guariglia DA, Tófolo LP, de Moraes SMF, Machado FA & Peres SB (2018). Peak velocity as an alternative method for training prescription in mice. Front Physiol; DOI: 10.3389/fphys.2018.00042.

Pollock RD, Carter S, Velloso CP, Duggal NA, Lord JM, Lazarus NR & Harridge SDR (2015). An investigation into the relationship between age and physiological function in highly active older adults. Journal of Physiology 593, 657–680.

Pollock RD, O’Brien KA, Daniels LJ, Nielsen KB, Rowlerson A, Duggal NA, Lazarus NR, Lord JM, Philp A & Harridge SDR (2018). Properties of the vastus lateralis muscle in relation to age and physiological function in master cyclists aged 55-79 years. Aging Cell; DOI: 10.1111/acel.12735.

Roman W, Martins JP, Carvalho FA, Voituriez R, Abella JVG, Santos NC, Cadot B, Way M & Gomes ER (2017). Myofibril contraction and crosslinking drive nuclear movement to the periphery of skeletal muscle. Nat Cell Biol 19, 1189–1201.

Ross JA et al. (2019). Impairments in contractility and cytoskeletal organisation cause nuclear defects in nemaline myopathy. Acta Neuropathol 138, 477–495.

Ross JA, Pearson A, Levy Y, Cardel B, Handschin C & Ochala J (2017). Exploring the Role of PGC-1α in Defining Nuclear Organisation in Skeletal Muscle Fibres. J Cell Physiol 232, 1270–1274.

Ross JA & Stroud MJ (2021). THE NUCLEUS: Mechanosensing in cardiac disease. International Journal of Biochemistry and Cell Biology 137, 106035.

Schindelin J, Arganda-Carreras I, Frise E, Kaynig V, Longair M, Pietzsch T, Preibisch S, Rueden C, Saalfeld S, Schmid B, Tinevez JY, White DJ, Hartenstein V, Eliceiri K, Tomancak P & Cardona A (2012). Fiji: An open-source platform for biological-image analysis. Nat Methods 9, 676–682.

Schoen I, Aires L, Ries J & Vogel V (2017). Nanoscale invaginations of the nuclear envelope: Shedding new light on wormholes with elusive function. Nucleus 8, 506–514.

Shah SB & Lieber RL (2003). Simultaneous Imaging and Functional Assessment of Cytoskeletal Protein Connections in Passively Loaded Single Muscle Cells. The Journal of Histochemistry & Cytochemistry 51, 19–29.

Shen Z, Lengyel M, Niethammer P & Os Lengyel M (2022). The yellow brick road to nuclear membrane mechanotransduction. APL Bioeng 6, 021501.

Shin JY & Worman HJ (2021). Molecular Pathology of Laminopathies. Annual Review of Pathology: Mechanisms of Disease 17, 159–180.

Shur NF, Creedon L, Skirrow S, Atherton PJ, MacDonald IA, Lund J & Greenhaff PL (2021). Age-related changes in muscle architecture and metabolism in humans: The likely contribution of physical inactivity to age-related functional decline. Ageing Res Rev 68, 101344.

Srivastava LK, Ju Z, Ghagre A & Ehrlicher AJ (2021). Spatial distribution of lamin A/C determines nuclear stiffness and stress-mediated deformation. J Cell Sci; DOI: 10.1242/jcs.248559.

Steger G (1998). An unbiased detector of curvilinear structures. IEEE Trans Pattern Anal Mach Intell 20, 113–125.

Stienen GJM (2000). Chronicle of skinned muscle fibres. Journal of Physiology 527, 1.

Stroud MJ (2018). Linker of nucleoskeleton and cytoskeleton complex proteins in cardiomyopathy. Biophys Rev 10, 1033–1051.

Stroud MJ, Feng W, Zhang J, Veevers J, Fang X, Gerace L & Chen J (2017). Nesprin 1α2 is essential for mouse postnatal viability and nuclear positioning in skeletal muscle. Journal of Cell Biology 216, 1915–1924.

Swift J, Ivanovska IL, Buxboim A, Harada T, Dingal PCDP, Pinter J, Pajerowski JD, Spinler KR, Shin JW, Tewari M, Rehfeldt F, Speicher DW & Discher DE (2013). Nuclear lamin-A scales with tissue stiffness and enhances matrix-directed differentiation. Science; DOI: 10.1126/SCIENCE.1240104.

Tajik A, Zhang Y, Wei F, Sun J, Jia Q, Zhou W, Singh R, Khanna N, Belmont AS & Wang N (2016). Transcription upregulation via force-induced direct stretching of chromatin. Nat Mater 15, 1287–1296.

Tan D, Yang H, Yuan Y, Bonnemann C, Chang X, Wang S, Wu Y, Wu X & Xiong H (2015). Phenotype-genotype analysis of Chinese patients with early-onset LMNA-related muscular dystrophy. PLoS One; DOI: 10.1371/journal.pone.0129699.

Wood DS, Zollman J, Reuben JP & Brandt PW (1975). Human skeletal muscle: Properties of the “chemically skinned” fiber. Science (1979) 187, 1075–1076.

Wroblewski AP, Amati F, Smiley MA, Goodpaster B & Wright V (2011). Chronic exercise preserves lean muscle mass in masters athletes. Physician and Sportsmedicine 39, 62.

York AG, Chandris P, Nogare DD, Head J, Wawrzusin P, Fischer RS, Chitnis A & Shroff H (2013). Instant super-resolution imaging in live cells and embryos via analog image processing. Nat Methods 10, 1122–1130.

Zhang J, Felder A, Liu Y, Guo LT, Lange S, Dalton ND, Gu Y, Peterson KL, Mizisin AP, Shelton GD, Lieber RL & Chen J (2010). Nesprin 1 is critical for nuclear positioning and anchorage. Hum Mol Genet 19, 329–341.

